# Asymmetric learning of dynamic spatial regularities in visual search: facilitation of anticipated target locations, no suppression of predictable distractor locations

**DOI:** 10.1101/2022.07.12.499748

**Authors:** Hao Yu, Fredrik Allenmark, Hermann J. Müller, Zhuanghua Shi

## Abstract

Static statistical regularities in the placement of targets and salient distractors within the search display can be learned and used to optimize attentional guidance. Whether statistical learning also extends to dynamic regularities governing the placement of targets and distractors on successive trials has been less investigated. Here, we applied the same dynamic cross-trial regularity (one-step shift of the critical item in clock-/counterclockwise direction) either to the target or a distractor, and additionally varied whether the distractor was defined in a different (color) or the same dimension (shape) as the target. We found robust learning of the predicted target location: processing of the target at this (vs. a random) location was facilitated. But we found no evidence of proactive suppression of the predictable distractor location. Facilitation of the anticipated target location was associated with explicit awareness of the dynamic regularity, whereas participants showed no awareness of the distractor regularity. We propose that this asymmetry arises because, owing to the target’s central role in the task set, its location is explicitly encoded in working memory, enabling the learning of dynamic regularities. In contrast, the distractor is not explicitly encoded; so, statistical learning of distractor locations is limited to static regularities.

**Public significance statement:** Can we learn the cross-trial dynamic regularity of a target or a task-irrelevant salient distractor (e.g., one-step shift of the critical item in clock-/counterclockwise direction) to boost search performance? The present study found robust learning of the predicted target location, but no evidence of proactive suppression of the predictable distractor location. Facilitation of the anticipated target location was associated with explicit awareness of the dynamic regularity. This asymmetry highlights the important role of the target-centered task set in the learning of dynamic regularities.

---

Our environment is extremely rich and complex, while our capacity for information processing is limited. The brain must prioritize information relevant to the task at hand, while resisting irrelevant information that might compete for our limited cognitive resources (Egeth & Yantis, 1997; Folk et al., 1992, 2002; Folk & Remington, 1998, 2008; Wolfe et al., 1989). Fortunately, rather than being random, our visual environment is highly structured. Accordingly, prior learning of environmental regularities and decisions could be useful for solving similar tasks. For example, it is easy to locate a sushi box from your familiar supermarket without being distracted much by other products, given that you know where the sushi boxes are. In the laboratory, this phenomenon has been systematically investigated in terms of so-called spatial ‘probability cueing effects’ (Geng & Behrmann, 2002, 2005). When a task-relevant target occurs with a high probability at one location, our attentional system can learn and effectively use this information for guiding search, facilitating target detection and response decisions (Druker & Anderson, 2010; Geng & Behrmann, 2002, 2005; Hoffmann & Kunde, 1999; Jiang et al., 2013; Shaw & Shaw, 1977).

Spatial probability cueing is not limited to prioritizing the target location. Rather, as has been shown in recent studies, high probability of a salient but task-irrelevant distractor appearing at a specific location or region can also be learned and used to de-prioritize the processing of such stimuli (e.g., Ferrante et al., 2018; Goschy et al., 2014; Leber et al., 2016; Sauter et al., 2018, 2019; B. Wang & Theeuwes, 2018a; Zhang et al., 2019). For example, Goschy and colleagues (2014) designed a visual search task that requires participants to search for a tilted bar amongst vertical bars and indicate whether the target bar had a gap at the top or the bottom. In half of the trials, a colored bar was shown with high probability (90%) in one half of the screen and with low probability (10%) in the other half. The ‘interference’ (i.e., the reaction time, RT, cost) engendered by a salient color distractor was greatly reduced if the distractor was presented in the high probability region, indicating that statistical learning of distractor locations can also boost search performance. In a control experiment, Goschy et al. further confirmed that the interference reduction is not merely owing to repetition of the distractor location across trials (which is more likely for likely distractor locations); rather, long-term statistical learning of likely distractor locations, and attendant suppression processes, contribute to the efficient search guidance.

Collectively, these studies have shown that observers can learn/exploit, from experience, the uneven spatial distributions of target and distractor in the search array over time, to minimize the interference generated by distractors and optimize target selection. However, whether statistical learning of target selection and distractor suppression are distinctive processes remains controversial. Some researchers argue that distractor suppression involves distinct processes to target selection (e.g., Noonan et al., 2016), while others suggest that attentional allocation by statistical learning is a result of a unitary mechanism: enhanced and, respectively, suppressed activities in spatial-attentional priority maps are just two sides of the same coin (Ferrante et al., 2018). It should be noted that statistical learning is not limited to the level of the spatial priority map. When the features or dimensions of the target and distractors are non-overlapped, selective attention can operate at the lower levels of features and dimensions in the functional architecture of the selective attention, as proposed by the Guided Search (Wolfe, 1994, 2021; Wolfe & Gray, 2007) and the Dimension-Weighting Account (DWA; Found & Müller, 1996; Liesefeld & Müller, 2019; Müller et al., 1995, 2003). For example, when color is never a target-defining dimension, such as in search for an orientation-defined target, the visual system boosts search efficiency by down-weighting signals from the (irrelevant) color dimension, and/or up-weighting signals from the (relevant) orientation dimension. The same would apply to feature selectivity: if a ‘red’ item is never a target, feature detectors tuned to red can be effectively down-weighted, and vice versa for detectors tuned to the critical target feature. Top-down dimension- or feature-based weighting processes, which operate prior to signal integration (across feature dimensions) by the priority map, are in principle spatially unspecific, that is, they operate in parallel and equally across the scene, and to what extent the weights can also be selectively tuned to particular display locations as a result of statistical learning (e.g., of distractor locations) is unclear (see Zhang et al., 2021 for a discussion). In any case, when dimension-based control is inapplicable (e.g., ignoring a salient intra-dimension distractor), spatially specific control of attentional selection based on statistical learning would have to operate at the level of the priority map, as has been shown in recent studies (Allenmark et al., 2019; Liesefeld & Müller, 2020).

The majority of the probability cueing studies demonstrating spatial statistical learning used a fixed uneven probability manipulation, such as one region/location having a higher occurrence of the target or distractor compared to the other region/locations (e.g., Geng & Behrmann, 2002, 2005; Goschy et al., 2014; Sauter et al., 2018; Shaw & Shaw, 1977). The implicit assumption is that statistical learning can modulate the activation pattern on the spatial priority map, by enhancing or suppressing specific locations/regions. However, the question remains whether or not such a modulation of selection priorities is stationary – adapting to a static spatial distribution of targets and distractors – or dynamic – adaptive to predictable changes in the distribution of targets and distractors. In a recent study, Li and Theeuwes (2020) introduced a dynamic cross-trial regularity to explore this question. In their paradigm, some target locations were predictably coupled across trials, such as a target occurring at the left-most (or respectively, the top) display location on trial *n* would invariably lead to the next target, on trial *n+1*, occurring at the rightmost (or respectively, the bottom) location (but not vice versa). Although apparently unbeknown to participants, (Li & Theeuwes, 2020) found this target regularity to nevertheless facilitate search and boost accuracy. In a more recent study, Wang et al. (2021) further explored whether such a flexibility would also characterize statistical learning, and attendant suppression, of distractor locations. In their adaptation of the ‘classical’ additional singleton search paradigm with a circular display arrangement, a salient color distractor ‘jumped’ by one location in either clockwise or, respectively, counterclockwise direction across consecutive trials, with 100% predictability. Wang et al. found that participants could relatively rapidly learn this cross-trial regularity to facilitate search, compared to a control group performing the task under conditions in which placement of the distractor across trials was random (i.e., the ‘regular’ group showed a reduced distractor interference relative to the distractor-absent baseline compared to the ‘random’ group). Note, though, that in their study the odd-one-out distractor color was either fixed (their Exp. 1) or randomly swapped between two colors (their Exp. 2), while the color of the (color-defined) distractor was *never* the color of the (shape-defined) target (white). Thus, as suggested by dimension-based accounts of distractor handling (Liesefeld & Müller, 2019), it would remain possible to globally, in a spatially non-specific manner, suppress feature contrast signals from the color dimension and so reduce their weight in the computation of the priority map. In fact, in Wang et al. (2021), the difference in interference between their ‘regular’ and the ‘random’ group was diminished towards the end of testing, suggesting that both groups were operating the same, spatially nonspecific suppression strategy (at least in the end); the reduced interference in the ‘regular’ group earlier on might then be explained by distractors occurring consistently across trials within the same (regularly moving) display region (of adjacent locations) increasing the learning rate compared to distractors occurring randomly in all display directions.

It has been shown that whether or not the target and distractor share the same defining dimension is a critical factor determining which strategy – priority-based or dimension-based suppression – is being adopted (Allenmark et al., 2019). Without color swapping between the distractor and the target (more precisely, between the distractor and *all* non-distractor items, the latter including the target), observers tend to adopt a dimension- (or feature-) based suppression strategy; with target-distractor color swapping, they develop priority-map-based suppression (Allenmark et al., 2019; Zhang et al., 2019). This echoes a similar idea proposed in the contingent capture hypothesis (Folk et al., 1992), namely, that the top-down attentional set for target-defining features determines which items are prioritized for selection: distractors can be effectively down-weighted only if they mismatch, rather than match, the features critical for discerning the target among the non-target items in the display – where the attentional control set influences signal coding below the level of the priority map, in a spatially nonspecific manner.

Thus, given that statistical learning of item locations for attentional prioritization can occur at multiple levels in the functional architecture of search guidance, whether dynamic enhancement and suppression are purely based on the predictive location of the target and, respectively, distractor remains elusive. To systematically investigate this, we devised the same cross-trial transitional probability structure for predictable target locations and, respectively, predictable distractor locations. In addition, we applied the same probability manipulation to distractors defined in the same and, respectively, a different dimension to the target (same- and different-dimension distractor, respectively) to establish whether (dynamic) statistical learning of distractor suppression would depend on the target-distractor dimensional relation. We hypothesized that if statistical learning of the predictable locations of the target and distractor are the ‘two sides of the same coin’, we should observe a similar pattern of dynamic spatial learning and attendant signal modulations – though in opposite directions: prioritization of target and de-prioritization of distractor signals – on the attentional priority map. By contrast, if dynamic modulation of spatial priorities by statistical learning is tied to the positive search goal, namely, to find some pre-specified target, we would expect to see a dissociation between dynamically predictable target locations (which should be learnable) and distractor location (which may not be learned, as they are only part of the negative task set). Specifically, in Experiment 1, we adopted the classic additional singleton paradigm and introduced cross-trial spatial regularities for the singleton color distractor (Experiment 1a) and, respectively, the singleton shape target (Experiment 1b, see Figure 1). The location of the critical item (either the target or the distractor) would move by one location across trials in one direction, either clockwise or counterclockwise (counterbalanced across participants) with a high probability (80%), or the opposite direction with a low probability (10%), or jump randomly to one of the non-adjacent locations (including location repetitions) (10%). Note that, in contrast to Wang et al. (2021), this implements a within-participant design (with the same participants performing both the regular and the random, baseline condition), avoiding spurious effects attributable to random group differences. To promote statistical learning taking place at the level of the priority map, we randomly swapped the target and the distractor color across trials, which previous research (Allenmark et al., 2019; Zhang et al., 2019) indicates limits learning at a level below the priority map (the level of specific features or feature dimensions). In Experiment 2, we used a letter search paradigm adopted from (Geng & Behrmann, 2002, 2005) and introduced a same-dimension distractor, rendering dimension-based distractor handling inapplicable. Again, the same cross-trial transitional probability structure was applied to the distractor (Experiment 2a) and the target (Experiment 2b). To preview the main results, we found robust statistical learning of dynamically predictable target locations, but failed to find any learning of dynamically predictable distractor locations.

**Figure 1.**
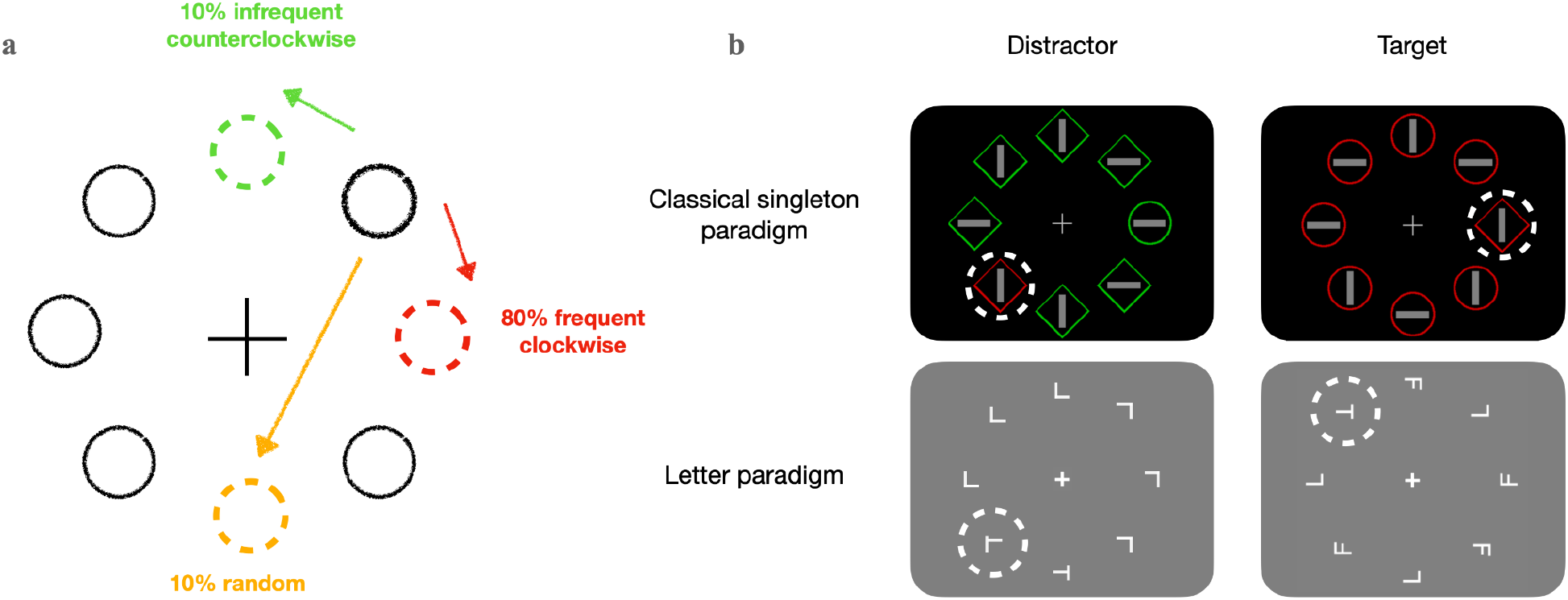
**(a)** Illustration of three cross-trial target- or, respectively, distractor-location transitions in Experiments 1 and 2. In each experiment, there were three types of the location change of the critical item (target or distractor) across consecutive trials: with 80% probability, the critical item would move to the adjacent location, either in clockwise or counterclockwise direction (here, indicated by the red dashed circle marking the frequent location). The direction was for a given participant and counterbalanced across participants. With 10% probability, the critical item would shift to the adjacent location in the opposite direction (indicated by the green dashed circle marking the infrequent location). On the remaining 10% of trials, the critical item would move randomly to any of the other locations, including re-appearing at the same location (indicated by the yellow dashed circle marking a random location). **(b)** Schematic illustration of four types of search display in which we implemented the cross-trial transitional regularity of the critical item (marked by white dashed circles, which were not presented in the experiments) to the left. The critical item was a color singleton distractor in Experiment 1a, the shape-defined target in Experiment 1b, the ‘T’-like distractor in Experiment 2a, and the ‘T’ target in Experiment 2b. As depicted in the upper panels, the target in Experiment 1 was an odd-one-out shape item (e.g., the circle at the 3-o’clock position in the upper left panel). In Experiment 2, the target was a left- or right-oriented T shape (e.g., at the 6-o’clock position in the bottom left panel).

## Experiment 1: Transitional regularity with the singleton paradigm

In Experiment 1, we applied the cross-trial transitional location regularity to the distractor (Experiment 1a) and the target (Experiment 1b) separately in a singleton target search paradigm with Experiment (a vs. b) as a between-subject factor.

### Method

#### Participants

24 healthy university students were recruited for Experiments 1a (mean age ± SD: 27.3 ± 4.2 years; age range: 21-39 years; 13 females) and 1b (mean age ± SD = 26.0 ± 3.2 years; age range: 21-33 years; 9 females) respectively. All participants reported normal or corrected-to-normal visual acuity. And all passed the Ishihara color test (Clark, 1924), ensuring they had normal color perception (especially for red and green). The participants can thus be regarded as representative of the standard population of healthy (young) adults.

The sample size was determined based on previous studies, in particular, Li and Theeuwes (2020), who had implemented a similar design introducing cross-trial regularities (for the target), with an effect size of *f* = 0.42 (average across all experiments). We conducted an a priori power analysis, with the effect size of *f* = 0.42, α = .05, and 98% power (1-β), which yielded a minimum sample size of *n* = 20 (G*Power 3.1; Faul et al., 2007). However, different from Li and Theeuwes, our study comprised both a singleton search and a letter search paradigm. Thus, to be on the safe side, we increased the sample size to 24 per tested group – an *n* that had also been used in another study with a similar design (Ferrante et al., 2018). All participants provided written informed consent prior to the experiment and were paid 9 Euro per hour or given correspondent course credit for their participation. This study was approved by the LMU Faculty of Pedagogics & Psychology Ethics Board. All data in Experiment 1 were collected in 2021.

#### Apparatus and Stimuli

The experiment was conducted in a sound-attenuated and moderately lit test room. Participants sat in front of the CRT display monitor, with a viewing distance of 60 cm. The search stimuli, presented at 1280 × 1024 pixels screen resolution and a refresh rate of 85 Hz, were generated by customized MATLAB R2019b (The Math-Works® Inc) code with Psychophysics Toolbox Version 3 (PTB-3) (Brainard, 1997).

As illustrated in Figure 1b, a search display consisted of eight items, each consisting of an outline shape (either diamond or circle) and a oriented bar (horizontal or vertical) inside it. The eight items were equidistantly arranged around an imaginary circle (radius 3.6° of visual angle). The diameter of the circle shapes was 1.4° of visual angle, the side length of the diamond shapes 1.9°, and the gray vertical or horizontal line inside the shapes 1.2° × 0.3°. Each display contained one singleton-shape target and seven non-targets. When a singleton distractor was present (replacing one of the non-targets), its color differed in color from the seven shapes shapes, being either green (CIE [Yxy]: [16.8, 0.306, 0.549]) among homogeneous red shapes (CIE [Yxy]: [11.6, 0.605, 0.336]), or red amongst homogeneous green shape. All search displays were presented on a black screen background (CIE [Yxy]: 1.72, 0.329, 0.265]), with a white fixation cross (0.76° × 0.76°; CIE [Yxy]: 79.7, 0.298, 0.298) in the center.

#### Design and procedure

A target, which was a shape-defined singleton (either a circle among diamonds or a diamond among circles, equally likely randomly assigned on each trial) was present on all trials. In order to realize a distractor-absent baseline in Experiment 1a without interrupting the structure of the cross-trial transitional probabilities of the distractor, we presented the singleton-distractor present and -absent trials in separate blocks. There were 16 blocks in Experiment 1a (4 singleton-distractor-absent blocks were randomly interleaved the other, singleton-present blocks). Each block consisted of 60 trials, yielding a total of 960 trials (240 distractor-absent trials and 720 distractor-present trials). Experiment 1b also consisted of 16 blocks, but without any singleton distractor. In both Experiments 1a and 1b, the target position was overall (across all trials) equally distributed among the eight possible locations, and participants had to respond to the orientation of the line inside the target as fast and accurately as possible.

Importantly, the placement of the critical item – the color singleton distractor in Experiment 1a, and the target singleton in Experiment 1b – across consecutive trials *n* and *n+1* was made predictable in a probabilistic manner. Specifically, in the majority of trials (80%), the location of the critical (distractor or target) item was shifted to an adjacent position in either clockwise or counterclockwise direction (with the main direction being fixed for a given participant, but counterbalanced across participants); hereafter, this will be referred to as the *frequent* condition. On another 10% of the trials, the position of the singleton distractor was shifted to the adjacent location in the opposite direction to the frequent condition (i.e., if the main direction was clockwise, the shift was counterclockwise, and vice versa) – the *infrequent* condition.^1^ And on the remaining 10% of the trials, the position of the critical item was randomly selected among the six remaining alternative locations (including repeated presentation at the same location) – the *random* condition. Of note, the statistical regularities were only assigned to the position of the singleton distractor or, respectively, the singleton target. Its color and shape varied randomly across trials. That is, the colors of the distractor and the target (as well as the other, non-distractor item) could randomly swap across trials – as in previous studies (e.g., Allenmark et al., 2019; Theeuwes, 1992), but different from Wang et al. (2021) design, in which the colors of the distractor were never the target color.

A trial started with a fixation cross presented in the center of the screen for 500 ms, followed by the search display (Figure 1b), which was shown until the participant gave a response. Participants were instructed to search for the shape-defined target and discriminate the orientation of the bar inside it by pressing the leftward- (‘horizontal’) or upward-pointing (‘vertical’) arrow on the keyboard with their right-hand index or middle fingers, respectively. If participants issued an incorrect response, a feedback display with the word “Error!” in the screen center was presented for 500 ms. The next trial started after an inter-trial interval of 500–750 ms. Between blocks, participants could take a break of a self-determined length.

At the end of the experiment, participants completed a post-experiment questionnaire in which they had to give two forced-choice responses: First, participants had to indicate whether or not they had noticed any regularity in the way Critical Items (CI: the target or distractor) had moved across trials. Next, they need to report the specific regularity of the movement, by choosing one of seven options for the most frequent type of movement (CI moved to opposite end of circle; CI moved one step clockwise; CI moved one step counterclockwise; CI moved two steps clockwise; CI moved two steps counterclockwise; CI moved three steps clockwise; CI moved three steps counterclockwise.)

#### Transparency and Openness

The experimental code, raw data, and data analyses of the present study are publicly available at: https://github.com/msenselab/asymmetric_statistical_learning. The pilot study (reported in the Appendix) was carried out in the winter semester 2018/19. Experiments 1 and 2 were conducted in 2021.

#### Bayesian analyses

Bayesian analyses of variance (ANOVAs) and associated post-hoc tests were carried out using JASP 0.15 (http://www.jasp-stats.org) with default settings. All Bayes factors for ANOVA main effects and interactions are inclusion Bayes factors calculated across matched models. Inclusion Bayes factors provide a measure of the extent to which the data support inclusion of a factor in the model. In more detail, inclusion Bayes factors compare models with a particular predictor to models that exclude that predictor: they indicate the amount of change from the prior inclusion odds (i.e., the ratio between the total prior probability for models that include a predictor and the prior probability for models that do not include it) to the posterior inclusion odds. Using inclusion Bayes factors calculated across matched models means that models that contain higher-order interactions involving the predictor of interest are excluded from the set of models on which the total prior and posterior odds are based.

### Results

#### Experiment 1a: transitional regularity of the distractor location

##### Error rates and Mean RTs

Trials with extreme RTs (slower than 2500 or faster than 200 ms) were excluded from further analysis (4.5% of trials). While the average error rate was overall low (4.7%), more errors occurred on distractor-present vs. -absent trials (5.4% vs. 3.4%), *t*(23)= 3.627, *p* = .001, *d_z_* = .74, with the error rates being comparable among the three (the frequent, infrequent, and random) distractor-location transition conditions, *F*(2, 46) = 1.002, *p* = .375, 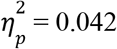, *BF*_incl_ = .286.

The mean (correct) RTs for the four distractor conditions (the distractor-absent baseline along with the frequent, infrequent, and random distractor-location transition conditions) are shown in Figure 2. As can be seen, the mean RT was faster in distractor-absent vs. distractor-present blocks, with the interference caused by distractor presence being significant, *t*(23) = 6.167, *p* < .001, *d_z_* = 1.26. Similar to the error-rate pattern, however, the RTs for the three cross-trial distractor-location transition conditions did not differ significantly among each other, *F*(2,46) = .168, *p* = .847, 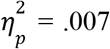, *BF*_incl_ = .134. That is, participants failed to learn the (frequent) cross-trial ‘movement’ of the location of the distractor to reduce its interference. This finding with a dynamic regularity of the distractor placement differs from that seen in ‘standard’ distractor-location probability-cueing paradigms, in which a fixed (stationary) frequent location/region of the distractor can be effectively learned to reduce distractor interference (Ferrante et al., 2018; Goschy et al., 2014).

**Figure 2.**
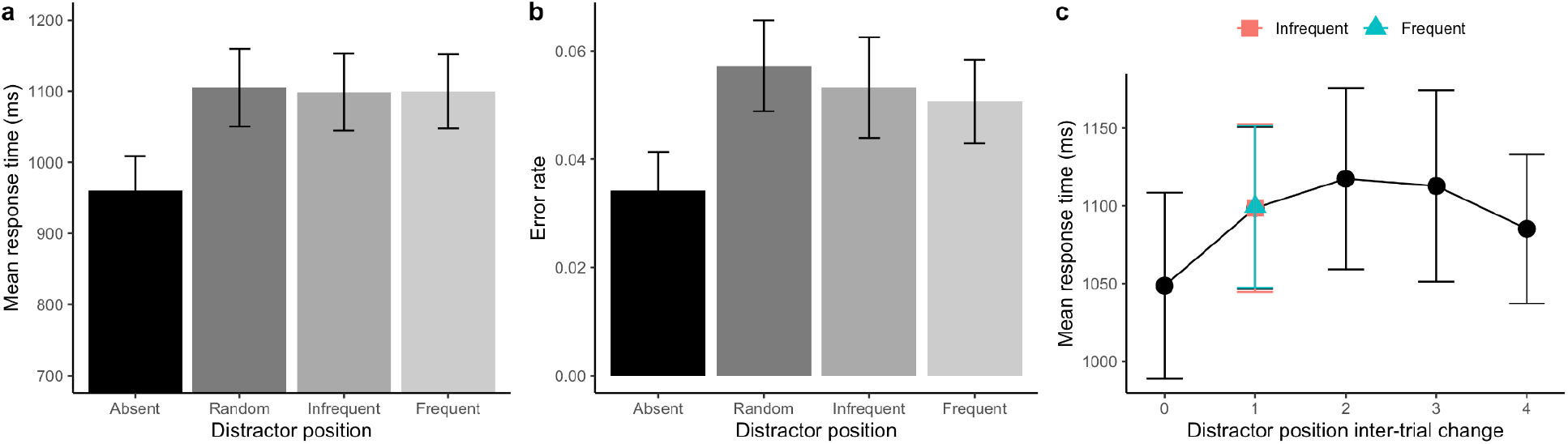
(**a**) Mean RTs and (**b**) Error rates, with associated standard errors, for Experiment 1a, separately for the distractor-absent baseline and the random, infrequent, and frequent cross-trial transitional distractor-location conditions. (**c**) Mean RT as a function of the inter-trial distractor distance (0 indicates the distractor repeated at the same location, while 1 denotes the distractor moved one position to its neighbor, including both the frequent and infrequent directions). Error bars represent one standard error of the mean.

It should be noted that the random cross-trial transition condition included exact location repetitions, which could potentially trigger (reactive) short-term inter-trial distractor-location suppression, facilitating search performance on such trials (see, e.g., the Supplementary in Sauter et al., 2018), for a detailed analysis of such effects). To examine for such inter-trial ‘negative-priming’ effects, we re-analyzed the mean RTs based on the inter-trial distractor distance (see Figure 2c, in which a distance of 0 denotes an inter-trial distractor-location repetition, while the distance of 1 includes all trials from the cross-trial frequent and infrequent distractor-location transition conditions, in an 8:1 ratio). Even though there was a numerical facilitation of 51 ms for distractor-location repetitions vs. the combined frequent and infrequent cross-trial transitions, *t*(23) = −1.922, *p* = .067, *d_z_* = −0.392, *BF*_10_ = 1.035, a repeated-measures ANOVA revealed the RTs to be (largely) comparable across the five inter-trial distances, *F*(4,92) = 1.970, *p* = .106, 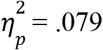, *BF*_incl_ = .445.

In standard (static) distractor-location probability-cueing paradigms (e.g., Goschy et al., 2014; B. Wang & Theeuwes, 2018a), it is often found that learned suppression of frequent distractor locations also impacts processing of the target when it occurs at such a location, evidenced by longer RTs to targets at frequent vs. infrequent distractor locations – typically assessed on distractor-absent trials. However, here, the distractor-absent trials were presented in separate (mini-)blocks from the distractor-present trials, so that it was not possible to estimate the target-location effect based on the distractor-absent trials. Thus, to examine for impeded target processing at the (dynamically predicted) frequent distractor location in our paradigm, we calculated the target-location effect based on our random distractor-location condition.^2^ A repeated-measures ANOVA with the factor Target Location (target occurring Frequent, Infrequent, and Random distractor location) failed to revealed any significant difference among the three target-location conditions (1104,1078, and 1099 ms for the Random, Infrequent, and Frequent locations, respectively), *F*(2,46) = 0.579, *p* = .564, 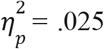, *BF*_incl_ = .179. This null-finding is consistent with the absence of a distractor-location effect (see Figure 3a).

**Figure 3.**
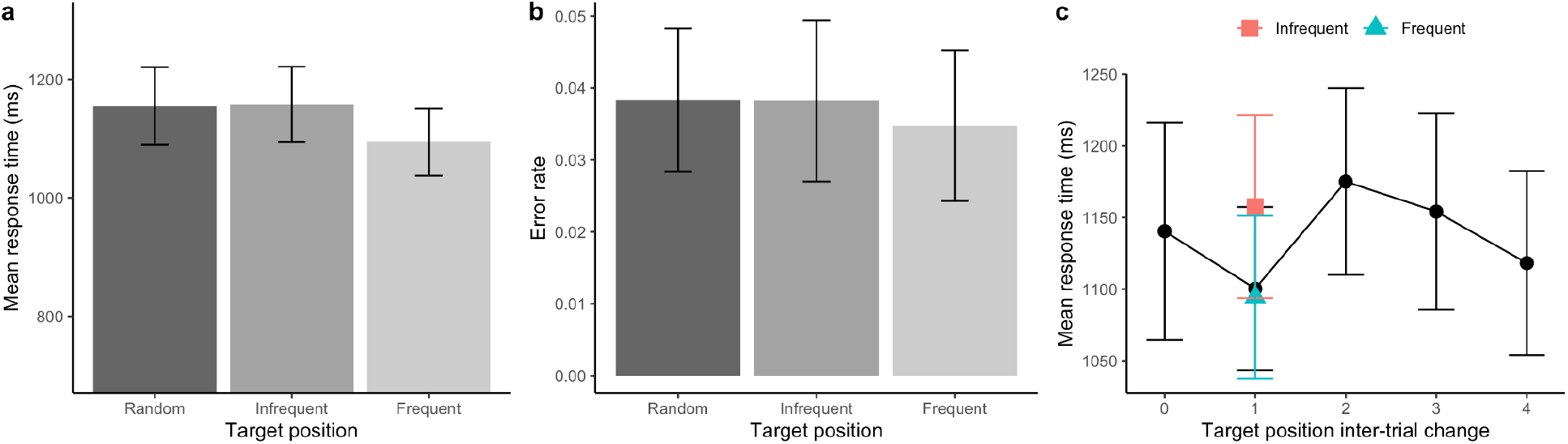
(**a**) Mean RTs and (**b**) Error rates, with associated standard errors, for Experiment 1b, separately for the random, infrequent, and frequent cross-trial transitional target location conditions. (**c**) Mean RT as a function of the inter-trial target distance (0 indicates the target repeated at the same location, while 1 denotes the target moved one position to its neighbor, including both the frequent and infrequent directions). Error bars represent one standard error of the mean.

##### Awareness test

Among the 24 participants, only three reported having noticed a regularity in the distractor movement. However, none of them was able to identify the specific regularity present in their search displays.

#### Experiment 1b: transitional regularity of the target location

##### Error rates and Mean RTs

Outliers RTs (slower than 2500 or faster than 200 ms, 6.0%) were again removed prior to further analysis. Similar to Experiment 1a, the error rates were generally low (3.5% of trials) and comparable across the three transitional target location conditions, *F*(2, 46) = .320, *p* = .728, 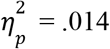, BF_incl_ = .162.

As depicted in Figure 3, the mean (correct) RTs were faster in the frequent cross-trial target-location transition condition relative to the infrequent and random conditions. A one-way repeated-measures ANOVA confirmed a significant Transition main effect, *F*(2,46) = 5.643, *p* = .006, 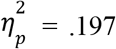. Post-host comparisons with Bonferroni-correction revealed the RTs to be faster in the frequent (1094 ms) vs. both the infrequent (1158 ms), *t*(23) = 2.970, *p* = .014, *d_z_* = 0.606, and random (1155 ms), *t*(23) = 2.845, *p* < .001, *d_z_* = 0.581, transition conditions, with comparable RTs between the latter two conditions, *t*(23) = −0.125, *p* = 1.000, *d_z_* = −0.009, *BF*_10_ = .219. This pattern indicates that participants were able to exploit the cross-trial transitional regularity of the target placement to facilitate search performance.

We also examined for short-term inter-trial positional-priming effects (e.g., Allenmark et al., 2019, 2021; Sauter et al., 2018) by comparing RT performance across the various inter-trial target distances (Figure 3c). A repeated-measures ANOVA, with the single factor of inter-trial Target Distance, failed to reveal revealed a significant Distance effect, *F*(4,92) = 1.753, *p* = .145, 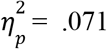, BF_incl_ = .329. The numerical facilitation, of 60 ms, for the inter-trial target distance of 1 vs. the target location repetition (distance 0) largely originated from the frequent cross-trial transition condition (which contributed 8 times more trials than the infrequent condition) (Figure 3a). This suggests that short-term inter-trial target location priming was not as strong as the dynamic, cross-trial probability cueing of the target location.

##### Awareness test

According to the questionnaire, 15 out of 24 participants reported noticing the regularity of the target movement, and ten of them indicated the right target movement direction. We classified those ten participants as the ‘aware’ group, and the other 14 participants as the ‘unaware’ group.

To examine for any differences between the two groups in statistical learning, we estimated the probability-cueing effect in terms of the RT difference between the infrequent and frequent transition conditions for individual participants. A positive probability cueing effect means that the mean RT is faster in the presence of the critical item at the frequent relative to the infrequent location, while a negative probability cueing effect indicates a reverse effect. Figure 4 plots the distribution of the probability-cueing effect for the two groups. The mean probability-cueing effects were 116 ms and 25 ms for the aware and unaware groups, respectively – with the effect being robust for the aware group, *t*(9) = 2.356, *p* = .043, *d_z_* = .745, but not for the unaware group, *t*(13) = 1.480, *p* = .163, *d_z_* = .396, BF_10_ = 0.660. Comparison between two groups revealed the probability-cueing effect was numerically higher for the aware vs. the unaware group, however it was not significant, *t*(22) = 1.970, *p* = .062, *d_z_* = .816, BF_10_= 1.445. This pattern suggests that becoming aware of the dynamic probabilistic change of the target location across trials helped participants to more effectively deploy visuo-spatial attention to the predicted target location.

**Figure 4.**
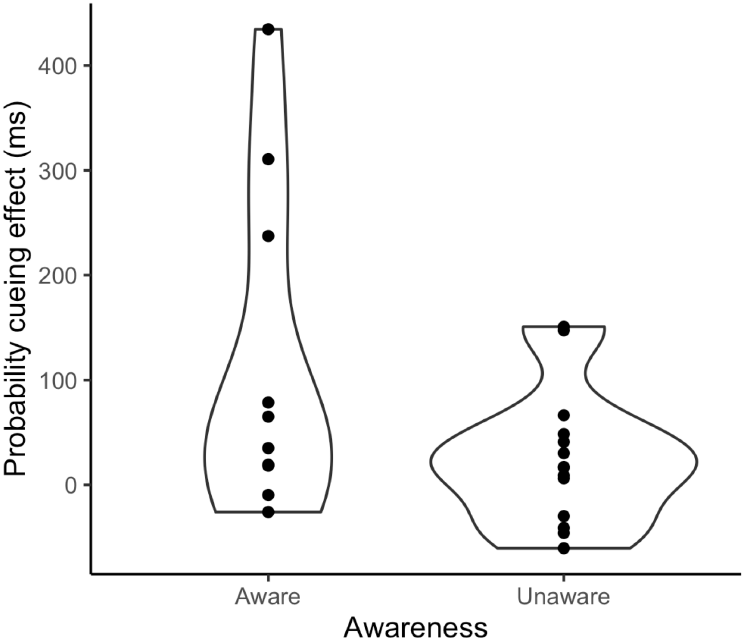
Violin plots of the probability-cueing effect (*RT_infrequent_* – *RT_frequent_*), separately for the aware and unaware groups of participants.

##### Comparison of probability-cueing effects between Experiments 1a and 1b

An independent-samples *t*-test comparing the probability-cueing effects between Experiment 1a and Experiment 1b (–1.1 ms vs. 63.3 ms) turned out to be significant: *t*(46) = −2.480, *p* = .017 (two-tailed), *d_z_* = −.716. In other words, participants could readily pick up the probabilistic change of the target position across trials and utilize it to enhance their search performance, whereas they found it hard to learn the same change of the distractor position across trials.

### Discussion

In Experiment 1, we manipulated the cross-trial transitional location probability of the singleton distractor (Experiment 1a) and the singleton target (Experiment 1b) in a standard (additional) singleton search paradigm. We found that the regularity of the cross-trial transition of the target location could be learned successfully to facilitate target search. In contrast, the dynamic regularity of the cross-trial distractor location had no significant effect on search performance and the Bayesian results supported the null hypothesis, even though the structure of the transitional probability manipulation was exactly the same for Experiments 1a as for Experiment 1b.

The ability to exploit the cross-trial regularity of the target placement to guide search is consistent with (Li & Theeuwes, 2020). In their study, however, the cross-trial regularity was 100% certain and relatively simple (either from the left- to the rightmost position, or from the top to the bottom location for half the participants, and in the reverse direction for the other half). Surprisingly, Li and Theeuwes reported that none of their participants had noticed this simple cross-trial regularity. The present experiment, by contrast, showed that awareness – that is, explicit learning – of the regularity boosted the dynamic target-location probability-cueing effect – suggesting that it reflects largely an endogenous, top-down-driven spatial-attentional orienting process (Posner, 1980).

In contrast to the facilitation by the transitional regularity of the target location, we failed to find any significant suppression of the dynamically predictable distractor location in Experiment 1a. This replicates the outcome of two pilot experiments with the same paradigm and a similar design (except that distractor-absent and present trials were presented in randomized order, rather than in mini-blocks, as in the present experiments; see Appendix 1 for details of the design and results). Although there is ample evidence that the probability of a fixed distractor location/region can be learned to suppress the salient distractor (Allenmark et al., 2019; Goschy et al., 2014; Sauter et al., 2018; B. Wang & Theeuwes, 2018b; Zhang et al., 2019), thus far there is only one study, by Wang et al. (2021), reporting that a regular (100% predictable) cross-trial change of the distractor location (clockwise or counterclockwise) could be implicitly (i.e., without awareness) learned to reduce the interference of the upcoming distractor. It should be noted, however, that in Wang et al. (2021), the colors of the distractor (single color in their Experiment 1, and two colors in their Experiment 2) were never the target color (the target was invariably white), and the differential distractor interference between their ‘random’ (baseline) group and their ‘regular’ group almost vanished towards the end of testing. Thus, it remains a possibility that the distractor-suppression strategy developed by their participants might involve dimension-based, or even feature-based, distractor filtering (Liesefeld & Müller, 2019), which operate below the level of the priority map. On this account, the cross-trial regularity would increase the rate with which (a spatially unspecific) dimension-based suppression strategy is acquired (compared to the ‘random’ baseline group), rather than fostering true learning, and attendant de-prioritization, of the dynamically predicted distractor location. In contrast to Wang et al. (2021), in our design, we randomly swapped the target and distractor colors across trials to make observers adopt a priority-map-based suppression strategy (Allenmark et al., 2019) – and failed to find any robust statistical learning of the cross-trial dynamics. The fact that responses were faster on trials on which the distractor appeared at the same (i.e., an unlikely) vs. the likely location (Figure 3b) suggests that distractor suppression was mainly driven by short-term inter-trial negative priming at the repeated (fixed) location, rather than the long-term learning of the dynamically predictable location.

One reason for our failure to find successful learning of the dynamic, cross-trial transitional distractor-location regularity could be that color was a task-irrelevant dimension, which can be down-weighted in general (Liesefeld & Müller, 2019; Müller et al., 1995), hampering the learning of statistical regularities in the placement of color singletons. The results of Experiment 1a suggest, by implication, that learning of dynamic distractor-location regularities requires participants to become aware of them, and becoming aware may be hampered if the regularity occurs in an effectively down-weighted, that is, (to-be-) ignored, stimulus dimension. It remains unclear whether the dynamic cross-trial regularity could be learned to suppress a predictable distractor location if dimension-based down-weighting is rendered impossible. In order to investigate this, we devised Experiment 2 with a singleton distractor defined within the target-defining dimension.

## Experiment 2: Transitional regularity with the letter paradigm

### Method

The setup was essentially the same as in Experiment 1, except the following differences.

#### Participants

24 healthy university students were recruited for Experiment 2a (mean age ± SD: 27.0 ± 4.02 years; age range: 19–37 years; 16 females) and 2b (mean age ± SD: 26.0 ± 5.3 years; age range: 19–39 years; 16 females) respectively. All participants were right-handed and had normal or corrected-to-normal vision. They all provided written informed consent prior to the experiment and were paid 9 Euro per hour or given correspondent course credit for their participation. All data in Experiment 2 were collected in 2021.

Due to the COVID pandemic, the planned Experiment 2b (transitional regularity of the target) had to be shifted online, through the online platform Pavlovia.

#### Apparatus and Stimuli

In the onsite Experiment 2a, the apparatus was identical to that in Experiment 1, while participants used their own computers in the online Experiment 2b. As depicted in Figure 1b (lower panel), the background of the search display was gray (CIE [Yxy]: [22.0, 0.304, 0.290]), and a white crosshair (0.48° × 0.48°, CIE [Yxy]: 79.7, 0.298, 0.298) was presented in the center of the display.

In Experiment 2a, the search items consisted of a “T”-shaped target presented among homogenous “L”-shaped non-targets; on distractor-present trials, one of the non-targets was replaced by a target-like distractor in which the intersected line in the “T” shape was not exactly bisected, but the intersection point was rather shifted slightly towards one end (i.e., the distractor was a cross between the target and non-target shapes). The search items were equidistantly arranged around a virtual circle 4° of visual angle in radius, and each item subtended 0.67° × 0.67° (CIE [Yxy]: 79.7, 0.298, 0.298); the small offset of the line junction in the distractor was 0.14° in size. The “T” target and “T”-like distractor were randomly rotated by 90° to either the left or the right from the upright orientation; the “L”-type non-targets were randomly rotated by either 0° or 180°.

In Experiment 2b, the search display consisted of eight shapes with a target “T” and non-targets “L” and “F”, with the items being randomly rotated 90° to either the left or the right. The “F”-shape non-targets were included to increase the difficulty of the task.^3^

#### Design and procedure

Experiment 2a consisted of 16 blocks (four distractor-absent, 12 distractor-present blocks), each of 60 trials (960 trials in total); and Experiment 2b consisted of 10 blocks, each of 84 trials (840 trials). The same structure of the cross-trial transitional regularity as in Experiment 1 (see Figure 1a) was applied to the distractor in Experiment 2a and the target in Experiment 2b. The task was to discriminate whether the base of the “T” target was pointing to left or right by pressing the F or J key on the keyboard as fast and as accurately as possible. Following the response, a feedback message (“correct” in green or “incorrect” in red) was shown for 500 ms.

At the end of the experiment, participants completed a post-experiment questionnaire in which they had to give two forced-choice responses: first, they had to indicate whether they had noticed any regularity in the “movement” of the critical item (the distractor in Experiment 2a, the target in Experiment 2b); next, participants were asked to select from seven options to indicate exactly how the critical item had moved.

### Results

#### Experiment 2a: transitional regularity of the distractor location

##### Error rates and Mean RTs

RT-outlier (1.2%) and response-error (2.8%) trials were relatively rare. Similar to Experiment 1a, errors were mainly made in distractor-present blocks (3.3%), rather than distractor-absent blocks (1.3%), *t*(23) = 4.413, *p* < .001, *d_z_* = −0.901. The error rates for the three cross-trial distractor-location transition conditions (frequent, infrequent, random; see Fig. 5b) were comparably low, *F*(2, 46) = 0.564, *p* = .573, 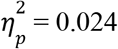, *BF*_incl_= .176.

**Figure 5.**
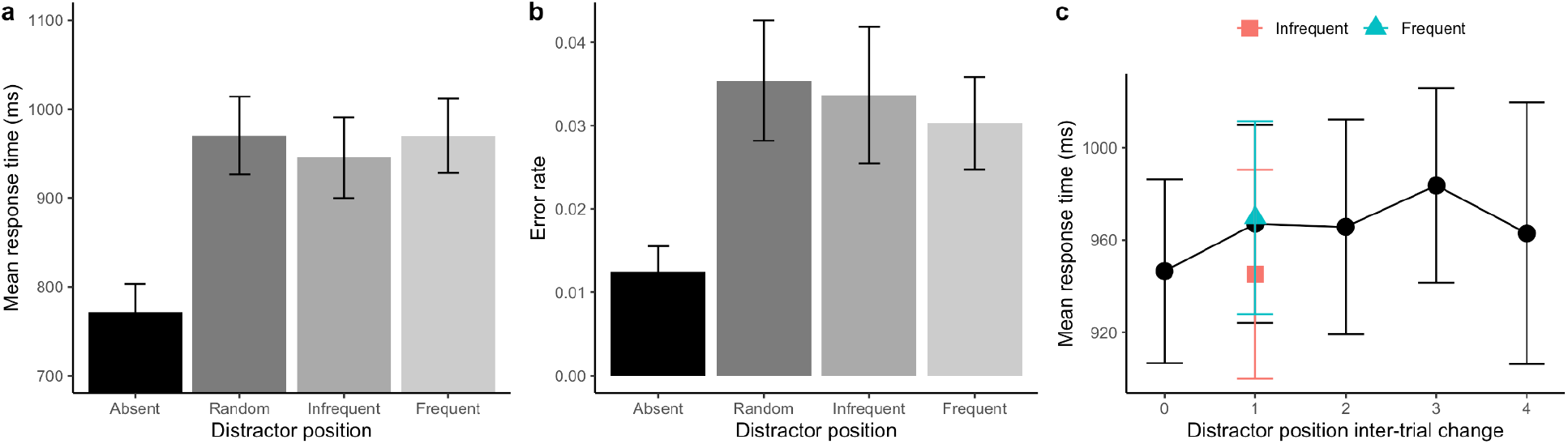
(**a**) Mean RTs and (**b**) Error rates, with associated standard errors, for Experiment 2a, separately for the distractor-absent baseline and the random, infrequent, and frequent cross-trial transitional distractor-location conditions. (**c**) Mean RT as a function of the inter-trial distractor distance (0 indicates the distractor repeated at the same location, while 1 denotes the distractor moved one position to its neighbor, including both the frequent and infrequent directions). Error bars represent one standard error of the mean.

Comparison of the (mean) correct RTs between the distractor-present condition(s) (967 ms) vs. the distractor-absent baseline (771 ms) revealed a significant distractor-interference effect (of 196 ms), *t*(23) = 13.118, *p* < .001, *d_z_* = 2.653 (Figure 5). A one-way repeated-measures ANOVA on the mean RTs in the three Transition conditions (frequent, infrequent, random transition; see Figure 5a) yielded a significant main effect, *F*(2,46) = 4.398, *p* = .018, 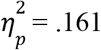. Surprisingly, post-hoc *t*-tests with Bonferroni correction revealed RTs to be faster for infrequent transitions (945 ms) compared to both frequent (970 ms), *t*(23) = −2.56, *p* = .042, *d* = −0.115, and random (970 ms), *t*(23) = −2.58, *p* = .039, *d_z_* = −0.116, transitions (there was no difference between the latter two conditions, *t*(23) = −0.027, *p* = 1.0, *d_z_* = −0.001, *BF*_10_ = .215).

Next, we again examined the mean RTs as a function of the inter-trial distractor distance (see Figure 5c). Similar to Experiment 1a, the response speed was comparable among the various distances, *F*(4,92) = 0.829, *p* = .510, 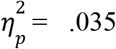, *BF*_incl_ = .093. There was a numerical RT benefit, of 21 ms, for distractor-location repetitions (distance 0) vs. cross-trial movements of the distractor to an adjacent position (distance 1, made up by frequent/infrequent transitions in an 8:1 ratio). Comparing the distractor-location repetition against the *infrequent* transition condition (which produced faster RTs than the frequent condition) failed to reveal the difference to be significant: 946 ms vs. 945 ms, *t*(23) = 0.065, *p* = 0.949, *d_z_* = 0.013, *BF*_10_ = .215. This suggests that the (unexpected) ‘facilitation’ effect observed with a distractor occurring at the infrequent location is related to lingering inter-trial negative priming, given that the distractor had appeared at that location on trial *n–1* (distractor-location-repeat trial) or on trial *n–2* (frequent transition trial, 80%). Nevertheless, we failed to find any evidence of distractor suppression at the most frequent location. This null finding is consistent with the results of Experiment 1a.

Additionally, a ANOVA examining for a Target-Location effect failed to reveal any evidence of RTs to a target occurring at the dynamically predicted (i.e., frequent) distractor location (957 ms) being slowed relative to targets appearing at a non-predicted (i.e., the infrequent or a random) distractor location (967 and 971 ms, respectively), *F*(2,46) = 0.166, *p* = .847, 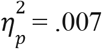, *BF*_incl_ = .132. This is consistent with the absence of reduced interference caused by distractors occurring at the predicted location (see Figure 5a).

##### Awareness test

Among the 24 participants, seven reported having noticed some regularities, but only three of them gave the correct response to the second question. Due to the small sample size of the aware group (*n*=3), statistical tests would not yield meaningfully interpretable results; accordingly, we refrained from further analysis.

#### Experiment 2b: transitional regularity of the target location

##### Error rates and Mean RTs

Again, there were only few outlier-RT (0.96%) and the response-error (3.2%) trials, and the error rates were comparable across the frequent, infrequent, and random cross-trial transitional target-location conditions, *F*(2, 46) = 1.223, *p* = .304, 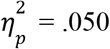, *BF*_incl_ = .291.

The mean RTs exhibited a systematic trend (see Figure 6a), being lowest for the random transition condition (1067 ms), intermediate for the infrequent condition (962 ms), and fastest for the frequent condition (893 ms). A one-way repeated-measures ANOVA confirmed the target-location Transition effect to be significant, *F*(2,46) = 30.63, *p* < .001, 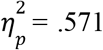. Post-host *t*-tests with Bonferroni correction revealed RTs in all conditions to differ significantly from each other, *t*s(23) > 3.09, *p*s < 0.01, *d_z_s* > 0.63.

**Figure 6.**
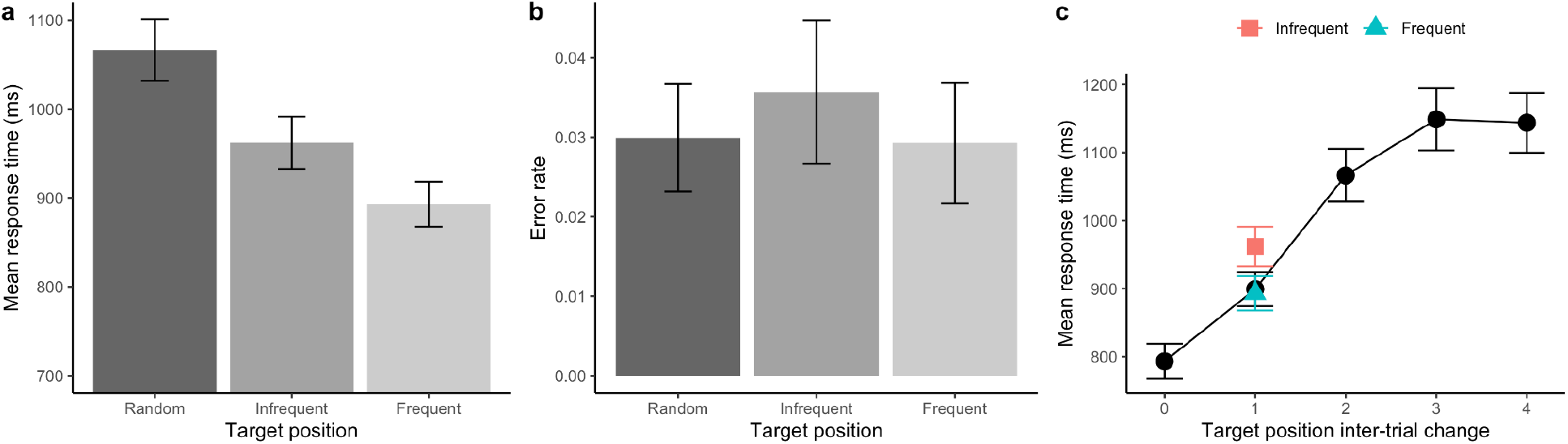
(**a**) Mean RTs and (**b**) Error rates, with associated standard errors, for Experiment 2b, separately for the random, infrequent, and frequent cross-trial transitional target-location conditions. (**c**) Mean RT as a function of the inter-trial target distance (0 indicates the target repeated at the same location, while 1 denotes the target moved one position to its neighbor, including both the frequent and infrequent directions). Error bars represent one standard error of the mean.

Figure 6c re-plots the mean RTs as a function of the inter-trial target distance. As can be seen, the RTs increased with increasing inter-trial target distance (up to distance 3) – a pattern quite unlike those seen in the other experiments. A repeated-measures ANOVA confirmed the Distance main effect to be significant, *F*(4,92) = 48.174, *p* < 0.001, 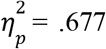. Post-hoc comparisons with Bonferroni correction revealed all but one comparisons to be significant, *ts*(23) > 3.29, *p*s < .014, *ds* > 0.59 (*d_z_*, non-significant comparison between distances 3 and 4, *t*(23) = 0.158, *p* = 1.0, *d_z_* = 0.029). The significantly faster RTs for distance 0 (exact target-location repetitions) vs. distance 1 (which is made up largely by the frequent target-location transitions) suggests that the inter-trial repetition facilitation outweighs the benefits deriving from the target moving to the predicted location.

##### Awareness test

Seven participants indicated being aware of a regularity in the placement of the target in the display, but only 4 of them correctly identified the regularity. Given the sample size was small for the aware group (*n* = 4), a statistical comparison with the unaware group would need to be interpreted with caution.

To visualize any awareness benefits for statistical learning of the dynamic target location, we calculated the probability-cueing effect (*RT_infrequent_* – *RT_frequent_*) for the individual participants, and divided them into the ‘aware’ and ‘unaware’ groups. Figure 7 shows the distribution of cueing effect for the two groups. Simple t-tests revealed the probability-cueing effect to be significant in the aware group, *t*(3) = 4.166, *p* = .025, *d_z_* = 2.083, but not in the unaware group, *t*(19) = 1.988, *p* = .061, *d_z_* = .444, *BF*_10_ = 1.184; further, the cueing effect was significantly larger in the aware than in the unaware group, (independent-samples) *t*(22)= 2.281, *p* = .033, *d_z_* = 1.249. In line with Experiment 1b, this pattern suggests that awareness of the predicted target location facilitated the guidance of attention to the dynamically predicted target location.

**Figure 7.**
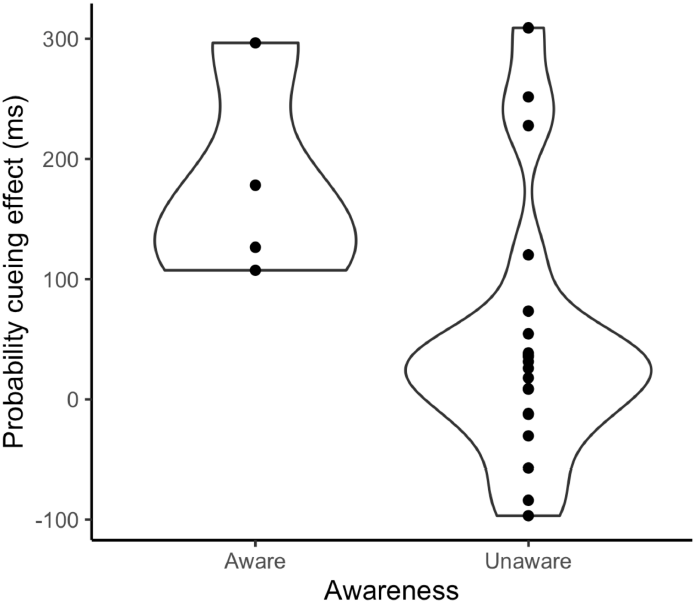
Violin plots of the probability-cueing effect (*RT_infrequent_* – *RT_frequent_*), separately for the aware and unaware groups.

##### Comparison of probability-cueing effects between Experiments 2a and 2b

An independent-samples *t*-test comparing the probability-cueing effects between Experiment 2a and Experiment 2b (–23.8 ms vs. 69.0 ms) turned out to be significant: *t*(46) = −3.689, *p* < .001 (two-tailed), *d_z_* = −1.065. This pattern is the same as in Experiment 1.

##### Comparisons across Experiments 1 and 2

We observed different probability-cueing effects between Experiment 1a and Experiment 1b, and between Experiment 2a and Experiment 2b. However, it is uncertain whether the results of Experiment 1 (singleton-search paradigm) and Experiment 2 (letter-search paradigm) are statistically different, that is, to which extent the task contributes to differential probability-cueing effects. To examine for any such contribution, we conducted a two-way factorial ANOVA on the mean RT probability-cueing effect with cross-trial Regularity (target, distractor) and Paradigm (singleton vs. letter search) as between-participant factors. The main effect of Regularity turned out significant, *F*(1,92) = 18.905, *p* < 0.001, 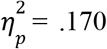, indicative of participants being able to pick up and utilize the cross-trial regularity in the positioning of the target, but not in the positioning of the distractor. Importantly, there was no evidence of the paradigm playing a role in this (Paradigm main effect, *F*(1,92) = .222, *p* = .638, 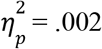, *BF*_incl_ = .237; Regularity × Paradigm interaction, *F*(1,92) = .617, *p* = .434, 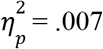, *BF*_incl_ = .368).

### Discussion

In Experiment 2, we changed the paradigm from the ‘classical’ (additional-) singleton paradigm in Experiment 1 to a paradigm that required search for a letter-type (shape) target in the possible presence of a letter-type (shape) distractor – that is, a distractor defined in the same dimension as the target: shape –, but we maintained the cross-trial transitional probability structure for the distractor location in Experiment 2a and for the target location in Experiment 2b. The result pattern turned out essentially the same as in Experiment 1: there was robust statistical learning of the dynamically predictable target location (RTs were significantly faster when the target to a location in the frequent, predictable vs. infrequent, non-predicted direction), but there was no evidence for learning of the distractor location (RTs were even slower when the distractor moved to a location in the frequent direction vs. the infrequent direction). The latter (non-) finding is at variance with a previous report by Wang et al. (2021).

Further, like Experiment 1b, Experiment 2b indicates that search performance is boosted by explicit awareness of the dynamic (probabilistic) target-location regularity – suggesting that the dynamic target-location cueing effect is based on endogenous control of visuo-spatial attention allocation. Further, in Experiment 2b, we found a strong inter-trial target-location repetition benefit – suggesting that search guidance is also modulated by a short-lived inter-trial priming effect, in addition to statistical learning of the dynamically changing target location.

## General Discussion

The present study was designed to investigate whether dynamic target-location enhancement and, respectively, distractor-location suppression purely based on the probabilistic cross-trial transitional regularities are possible. We conducted two experiments with the same cross-trial transitional regularity (80% likely one-step clockwise or counterclockwise ‘movement’) of the critical item: either the search target or a task-irrelevant distractor. In addition, we manipulated how the distractor was defined in relation to the target: in a different dimension (color; Experiment 1) or the same dimension (shape; Experiment 2). We found robust dynamic search guidance when the target location shifted predictably across consecutive trials. In contrast, we failed to find any evidence of reduced distractor interference when the distractor location shifted predictably across trials, not even when the distractor was defined in the same dimension as the target. Facilitated processing of the target at the predicted location appeared to be associated with conscious awareness of the dynamic regularity: only those participants who explicitly recognized the regularity exhibited a robust facilitation effect. In contrast, there was no evidence of participants becoming aware of the regularity in the distractor item, not even it was defined in the same dimension as the target.

The statistical learning of the predicted target location that we observed in both experiments broadly consistent with the probability cueing of the target location reported in the literature (Geng & Behrmann, 2002, 2005; Shaw & Shaw, 1977). For example, manipulating the likelihood of target presentation unevenly between the left and right sides of the display (80% vs. 20%), Geng and Behrmann (2002) found search to be facilitated when the target actually appeared within the more probable region. Of note, though, most of the previous target-location probability-cueing studies used a spatially fixed (or stationary) uneven probability manipulation (either location- or region-based), finding that search guidance can remarkably adapt to these environmental statistics to boost performance. A recent study, by Li and Theeuwes (2020), showed that this adaptability also extends to dynamic location manipulations: when the target on trial *n-1* (appearing, e.g., at the leftmost display location) predicted the location of the target on trial *n* (in the example, the right-most location) with 100% certainty, participants were also able to learn this cross-trial regularity to facilitate search performance. Here, we showed that a dynamic cross-trial regularity can also be learned when it is probabilistic in nature. Similar to earlier studies (e.g., Geng & Behrmann, 2002), we implemented an uneven cross-trial transitional probability structure (80% for cross-trial frequent, 10% for infrequent, and 10% for random transitions) and showed that participants could learn this probabilistic regularity and use it to facilitate target detection. It is important to note that, in our study, the global probability of the target occurrence remained equal across all possible locations – only the cross-trial transitional probability differed in the direction of the target movement (clockwise or counterclockwise). This suggests that the search-guidance system can learn and adapt to both fixed and dynamic probability structures that govern where the target appears, and modify the weights on the attentional-priority map accordingly.

In contrast to robust cross-trial dynamic probability-cueing of the target location, we failed to find any evidence that participants were able to learn the same dynamic probability structure when this was applied to predict distractor location. This is different from the many studies with a fixed uneven distribution of the distractor, which have collectively shown that display locations/regions with a high probability of distractor occurrence can be effectively de-prioritized to reduce the interference caused by the irrelevant pop-out stimulus (Ferrante et al., 2018; Goschy et al., 2014; Leber et al., 2016; Sauter et al., 2018, 2019; B. Wang & Theeuwes, 2018a; Zhang et al., 2019). For instance, likely distractor locations may be proactively suppressed – that is, some ‘no-go’ tag may be placed on them – on the attentional priority map (e.g., Ferrante et al., 2018), dampening the built-up of the priority signal at such locations. Thus, while ‘no-go’ tagging of *fixed* likely distractor locations is a feasible strategy for the search-guidance system to reduce attentional capture, our findings suggest that ‘no-go’ tagging of *dynamically* predictable distractor locations is not possible – at least not with the same dynamic probability structure (and the same number of learning trials) that we used for the target location.

Our non-finding also appears to be at variance with Wang et al. (2021), who reported that a very similar cross-trial transitional regularity of the distractor (clockwise or counterclockwise movement by one step). Importantly, however, there are several key differences between their study and ours. First, the regularity they implemented was deterministic (100%), rather than probabilistic (our structure predicted the distractor location with 80% probability). Whether this is a critical difference is, ultimately, an empirical matter. However, there is evidence from reward-association learning of ‘incentive salience’ that probabilistic regimens are more effective than deterministic regimens (e.g., Cho & Cho, 2021; Sali et al., 2014). Also, our participants had no problem learning exactly the same deterministic structure in relation to the target location. Another key difference in Wang et al.’s (2021) study from ours is that the color of the (color-defined) distractor was distinct from the fixed target color in their experiments (even in their Experiment 2, in which the distractor could appear in two possible colors). In our color-distractor experiment (Experiment 1a), we purposely implemented random swapping, across trials, of the distractor and target (i.e., more generally, the non-distractor) colors to reduce possible dimension- (or feature-based) distractor suppression which operates in a spatially non-specific manner. As shown in our previous studies (Allenmark et al., 2019; Zhang et al., 2019), whether participants adopt dimension-/feature-based or a priority-map-based suppression much depends on the overlapping of the distractor and target features: with color swapping between target and distractor, participants tend to adopt a priority-map-based suppression strategy; without color swapping, they are likely to develop a dimension-based strategy (reducing the weight of color signals in priority computations). This is consistent with Gaspelin and Luck (2018), who also reported that with random trial-by-trial swapping of the singleton and non-singleton colors in the search display, (post-display) probe suppression effects (i.e., reduced report accuracy for probes at the distractor location vs. averaged non-distractor locations) were completely eliminated. Similarly, in an eye-movement experiment with color swapping between distractors and targets, Gaspelin and Luck (2018) found the first saccade (after display onset) to be more likely to be directed to the singleton distractor than to the average of the other non-target items, in line with color swapping reducing lower-level distractor suppression. Accordingly, participants in Wang et al.’s (2021) study might have adopted a spatially unspecific strategy of suppressing the color dimension in both the condition with the ‘regular’ structure (performed by one group of participants) and the baseline condition with the ‘random’ structure (performed by another group). Some evidence of this is provided by the fact that, while distractor interference generally (i.e., in both groups) decreased across the ten trial blocks of their experiments, there remained little difference in interference between the ‘regular’ and the ‘random’ groups by the end of testing. That is, the ‘regular’ group only showed a faster rate of interference reduction over the course of the experiments – which could mean that learning to suppress the irrelevant color dimension works more efficiently when the distractor occurs in a contiguous display region across trials (such as in a stepwise movement of the distractor from one location to the next) compared to when it crisscrosses the display in a random manner.^4^ Whatever the precise reason(s) for the discrepant findings between the Wang et al. (2021) and our Experiment 1a, it remains (at least) that it is difficult to demonstrate successful learning of a dynamic probabilistic regularity with regard to the locations of (color-defined) distractor, but not with regard to the locations of color-defined targets.

Of note, despite attempting to make dimension-based distractor suppression hard in Experiment 1a (by introducing random color swapping), we could not rule out that participants nevertheless developed such a strategy – given that color was completely irrelevant to solving the task to find a shape-defined target. If so, color feature-contrast signals would have been globally suppressed (at least to some extent), impeding statistical learning of the dynamically predictable location of the *color* distractor. As a consequence, statistical learning of the predictive location of the *color* distractor might have been reduced or diminished – and this is why we found no distractor-location cueing effect in Experiment 1a. Therefore, in Experiment 2a, we introduced a distractor defined within the same dimension as the target: a shape-defined distractor. Given that participants had to set themselves for a shape target, they could not globally ignore the shape dimension, as this would have conflicted with the task goal. As expected (e.g., Geng & Behrmann, 2002, 2005; Goschy et al., 2014; Sauter et al., 2018; Shaw & Shaw, 1977), the shape distractor caused massive interference, of 185 ms relative to the distractor-absent baseline – which is likely due to ‘overt’ attentional capture, involving a first eye movement to the distractor on most trials before attentional/oculomotor disengagement and re-orientation to the target (see Sauter et al., 2021). In other words, the distractor was expressly processed as a ‘wrongly’ selected item – and yet, participants failed to learn its dynamic cross-trial transitional regularity. In contrast, when implemented in the target placement (Experiment 2b), the same regularity again produced a dynamic facilitation effect, as in Experiment 1b.

Thus, the question remains why dynamic suppression of predictable distractor locations is so hard (if not impossible), whereas dynamic facilitation of predicted target locations is established easily. A clue to answering this question is provided by the ‘awareness’ results. In both Experiments 1b and 2b, participants became substantially aware of the dynamic target regularity, and those of them who correctly selected the right regularity (out of seven alternatives) in the awareness test showed a significantly larger facilitation effect than the ‘unaware’ participants. Pooling the two experiments together for improved statistical power to compare the facilitation effects between the aware and unaware groups revealed a robust benefit of explicit awareness, *t*(46) = 2.811, *p* = .007, *d_z_* = 0.893. This is not to say that the dynamic target regularity cannot be implicitly learned. In fact, the pooled ‘unaware’ group also exhibited a significant facilitation effect, of 38 ms (*t*(33) = 2.456, *p* = .020, *d_z_* = 0.421), which however is only a small fraction of the 134-ms effect exhibited by the aware group (134 ms).^5^ The fact that explicit awareness greatly boosted the dynamic facilitation effect suggests that participants did develop a dynamic top-down set to prioritize the next location in the regular (clockwise or counterclockwise) direction in anticipation of the next target occurring there (endogenous orienting in Posner, 1980 terms). We suggest that they did develop such an anticipatory top-down set because the target is the central item in the task set: observers have to set up a target template in working memory and compare any selected item against this template, and then deselect it if there is a mismatch or proceed to extracting the response-relevant feature and select the appropriate response if there is a match. Given the central place of the target in the task set, even seemingly irrelevant ‘features’ such as its location may be explicitly encoded, providing the basis for recognizing the regularity in the placement/movement of the target across consecutive trials. In contrast, if a distractor is mistakenly selected, it only needs to be rejected as a non-target item, that is, as not matching the target template; in other words, there is no need to process the distractor for, and explicitly represent, any featural information about the distractor, including its location. As a result, there is no explicit learning of higher-order dynamic statistical regularities in the placement of the distractor.

However, there is implicit learning of static statistical regularities, that is, of a fixed display location or region being more likely to contain a distractor than other locations – as evidenced by a plethora of recent demonstrations of distractor-location probability-cueing effects in the absence of conscious awareness of bias in the distractor distribution. We have recently shown that these static cueing effects depend purely on the the local distractor probability (Allenmark et al., 2022), and that the frequency with which distractors occur at a particular location modulates the responsivity of neurons in early (i.e., retinotopic) visual cortex areas, from V1 to V4 – with higher frequency rendering a stronger down-modulation (Zhang et al., 2021). Also, we proposed that the ‘tuning’ signal for the down-modulation of entry-level cortical feature coding is provided by the ‘rejection’ (or reactive suppression) of a particular location when a selected (distractor) item at this location produced a mis-match decision: the more often this happens for a particular locations, the lower the responsivity of V1–V4 neurons with corresponding receptive fields becomes – which naturally explains the static distractor-location probability-cueing effect. That is, this is an essentially static mechanism (top-down inhibiting the current distractor location, so as to disengage attention and re-deploy it to the target location), which does not require conscious knowledge of the distractor location to work. Given this, it is hard to see how it could be ‘dynamicized’ to track a distractor that changes position predictably, according to some higher-order rule, across trials.

In contrast, a dynamically predictable target location can be tracked successfully if the rule is explicitly (consciously) represented in working memory, as part of the task set. This rule can then be applied to flexibly prioritize a given next location, given the current target location, perhaps by top-down pre-activating the anticipated location on the attentional priority map. Further, neuroscientific work is necessary to examine the brain mechanisms underlying dynamic target-location prediction, though these are likely to involve the frontoparietal attention network: a richly interconnected network linking the intraparietal sulcus (IPS), the inferior parietal lobe (IPL), and dorsal premotor cortex (PMC), including the frontal eye field (FEF). According to Ptak’s (2012) model of this network, the posterior parietal cortex has functional characteristics that point to a central role of this region in the computation of a feature- and dimension-independent attentional-priority map. “Feature maps computed in the sensory cortex and current behavioral goals as well as abstract representations of associated actions (action templates) generated in the prefrontal and premotor cortex (PMC) feed into the parietal priority map. The dorsolateral prefrontal cortex (DLPFC) maintains behavioral goals in working memory and protects them from distracting information. The inferior parietal lobe (IPL) initiates shift of attention and maintains attention on the relevant stimulus” (Ptak, 2012, p. 512). Given this, it is conceivable that *dynamic* spatial expectations originating in the DLPFC and PMC can also be integrated in the priority map.

## Conclusion

The present study investigated statistical learning of the same dynamic, cross-trial probabilistic regularity of the target and (additional-singleton) distractor location in visual search. In several experiments, we consistently found robust facilitation of the dynamically predictable target location, but no suppression of the dynamically predictable location of the distractor (the latter being at variance with another report in the literature). While none of the participants noticed the cross-trial regularity of the distractor, one third of the participants correctly selected the cross-trial target regularity in a post-search explicit-recognition test; further, awareness of the target regularity greatly (by a factor of 4) enhanced cross-trial cueing of the target location. We propose that this asymmetry, in the dynamic cueing and awareness effects, arises because the target occupies a central place in the task and so is explicitly encoded in working memory for template matching and extraction of the response critical feature; as a result, the dynamic cross-trial change in its location is also registered and can be used to top-down prioritize the upcoming target location. In contrast, the distractor is not an explicit part of the task set; (e.g., no distractor template needs to be set up in working memory to reject a distractor that captured attention), and so statistical learning of the distractor location is only static, limited to its current position and how frequently it occurs at this position.

## Appendix 1: Analysis of pilot experiment data

In two pilot experiments, intended to investigate whether observers can learn to suppress a predictable distractor location based on dynamical regularities in the trial-to-trial movement of the distractor, we obtained similar results to Experiments A1a and 2a. These experiments used essentially the same paradigm (stimuli and task) as in Experiment 1a, with a few minor differences. The distractor was more likely to move in one, ‘frequent’ direction, either clockwise or counterclockwise (counterbalanced across participants), but on the remaining trials it moved randomly to any other location (i.e., unlike Experiments 1a, movement in the direction opposite to the frequent direction, the ‘infrequent’ direction was no more likely than movement to any other location). In both experiments, a distractor was present on overall 66% of trials, but (unlike Experiment 1a) distractor-absent trials were randomly interleaved with distractor-present trials. When one or more distractor-absent trials occurred in-between two distractor-present trials, the ‘frequent’ distractor location on the second of the two interrupted distractor-present trials was defined in two different ways: for half of the participants the frequent location was one step in the frequent direction compared to the last distractor-present trial, while for the other half the frequent distractor location was the location where the distractor would have been if a distractor had been present and moved in the frequent direction on each trial (e.g., with two distractor-absent trials in-between two distractor-present trials, the frequent location on the second distractor-present trial was three steps in the frequent direction from the distractor location on the first distractor-present trial). In pilot Experiment 1, 65% of distractors occurred at the frequent location; pilot Experiment 2, this probability was increased to 85%. We tested 14 participants in Experiment 1 and 16 in Experiment 2.

Figure A1 shows the mean response times and error rates in both pilot experiments in the distractor absent condition, as well as for trials with distractors appearing in the frequent location and random locations. Distractor interference – the difference in RT between distractor-present and distractor-absent RTs – was significantly greater than zero in both experiments (Exp. 1:118 ms, *t*(13) = 6.40, *p* < .001; Exp. 2: 117 ms, *t*(15) = 7.95, *p* < .001). However, there was no significant difference between RTs when a distractor occurred in the frequent location compared to a random location in either experiment (Exp. 1: 2 ms, *t*(13) = 0.15, *p* = .88; Exp 2: −12 ms, *t*(15) = −1.30, *p* = .21). Since the predictability of the frequent distractor location could be reduced by interruptions by distractor absent trials we also checked whether there was a difference between RTs when a distractor occurred in the frequent location compared to a random location when considering only pairs of consecutive distractor present trials but there was still no significant difference (Exp 1: 5 ms, *t*(13) = 0.36, *p* = .72; Exp 2: −12 ms, *t*(15) = −1.15, *p* = .27).

**Figure A1.**
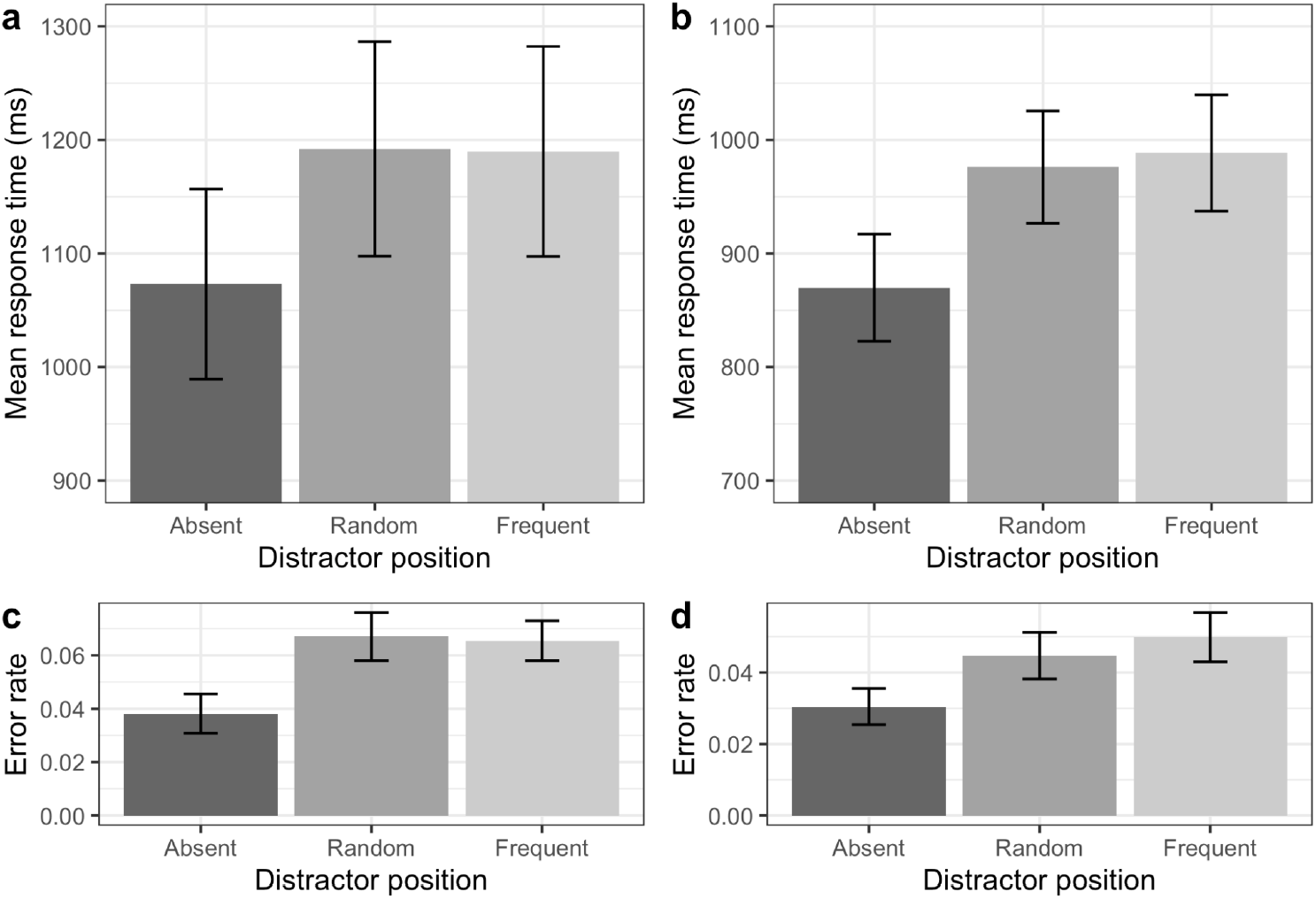
Mean response times (a, b) and error rates (c, d) on distractor absent trials, trials with a distractor in the (dynamic) frequent distractor location and on trials with a distractor in a random location in pilot experiment 1 (a, c) and pilot experiment 2 (b, d). Error bars represent one standard error of the mean.

## Footnotes

1 This condition was introduced to allow us to compare two conditions with the same inter-trial distance (movement of the critical item by one step) but different probability. With only the frequent and random conditions, we would have had too few trials on which the critical item moved the same distance in the random as in the frequent condition (but in the opposite direction). Figure shown in 6c, comparing conditions with the equal distance turned out ‘diagnostic’ at least in one of the experiments.

2 Note that Wang et al. (2021) could not examine for such an effect because, in their design, the distractor appeared with 100% certainty at the location one step ahead of the current location in a given direction. That is, a target could never occur at this perfectly predicted location.

3 We considered this necessary in Experiment 2 but not in Experiment 1, because in the latter the random trial-to-trial swapping of the target and non-target shape (circle among diamonds and diamond among circles) ensured sufficient task difficulty. In contrast, searching for a “T” target among homogeneous “L”-shape non-targets in the absence of any distractor would have made the task easier. Thus, we introduced an element of non-target heterogeneity in Experiment 2 to avoid a possible ceiling effect – attributable to the target being invariably the first item to summon attention, which may have left insufficient ‘room for improvement’ by learning of the dynamic target-location regularity.

4 Alternatively, their result pattern might also be explained by reactive suppression placed post-capture on the distractor location, in order to disengage attention and re-orient it to the target. If reactive suppression is somewhat fuzzy, affecting adjacent locations, and if it is carried over across trials, it would, on average, have a greater impact with the regular movement of the distractor to an adjacent location, as compared to the random placement. Wang et al. (2021) argued against this possibility based on an analysis of inter-trial negative priming effects in their ‘random’ condition (with a different group of participants). They could not examine for such effects directly in the ‘regular’ group, because the dynamic distractor regularity was deterministic. Note in this context that our ‘random’ baseline was a within-participant, rather than a between-participant, manipulation, ruling out confounding of effects by spurious group differences.

5 Interestingly, apparently none of the participants in Li and Theeuwes (2020) reported noticing the cross-trial target regularity that they had encountered during the search (where a target at, say, the leftmost location predicted with certainty that the next target would appear at the rightmost location).

## Notes

**Author note**, This study was supported by German Science Foundation (DFG) research grants MU773/16-2, awarded to HJM and ZS, and MU773/14-2, awarded to HJM. Data and analysis code of the present study are available at: https://github.com/msenselab/asymmetric_statistical_learning.

### Competing Interest Statement

The authors have declared no competing interest.

